# SELECT-seq allows Pre-Sequencing Enrichment of SNP Edits in One-Pot Single-Cell Whole-Transcriptome Sequencing

**DOI:** 10.64898/2026.07.07.736965

**Authors:** Sho Iwama, Daniel Gitterman, Timothy Brendler-Spaeth, Andrew Waters, Holly Robertson, Magdalena E. Strauss, Dave Adams, Sarah E. Cooper, Qianxin Wu, Andrew R. Bassett

**Affiliations:** Wellcome Sanger Institute, Wellcome Genome Campus, Hinxton, CB10 1SA, UK; University of Exeter, Exeter, Exeter, EX4 4QD, UK

## Abstract

Advances in high-throughput sequencing have associated millions of putative genetic variants with disease. However, scalable experimental methods to establish causal relationships between genetic variants and downstream transcriptional outcomes remain a major challenge. Single-cell methods that integrate genotyping with transcriptomic profiling provide a way to address this, but do not enable pre-sequencing enrichment of correctly edited cells, limiting scale.

We present SELECT-seq (SNP Enrichment Leveraging Cas12a Targeting), a rapid method that allows SNP-specific PCR amplification and Cas12a-mediated fluorescence detection simultaneously with whole-transcriptome amplification. This one-pot workflow enables identification and enrichment of SNP-bearing single cells, making a rapid and scalable methodology for analysis of genotype-phenotype linkage avoiding laborious single cell cloning steps.

As a proof of principle we show that SELECT-seq distinguishes U-2 OS and T-47D cell lines based on a *PIK3CA* (NM_006218.4:c.3463A>G) mutation while preserving transcriptome integrity. It physically enriches a rare NRF2 T80K (NM_006164.5:c.390C>A) mutant cells (6.7%) from a prime-edited pool, achieving 86% genotype accuracy, and shows 87.5% directional concordance in the transcriptomic effects compared with a clonal NRF2 T80K cell line. SELECT-seq thus provides a rapid, scalable and widely accessible approach for mapping genotype–phenotype relationships at single-cell resolution.

## Introduction

Large-scale sequencing efforts such as genome-wide association studies (GWAS, https://www.ebi.ac.uk/gwas/), genome and exome sequencing (https://www.ncbi.nlm.nih.gov/clinvar/) have catalogued millions of human genetic variants that have been associated with disease, with single-nucleotide polymorphisms (SNPs) representing the most abundant class. Despite this, the functional consequences of most variants remain poorly understood, with around 35-50% of variants in ClinVar being variants of uncertain significance (VUS)(1).

Base editing, prime editing and homology-directed repair (HDR) enable the introduction of defined variants such as SNPs to understand their function. However, even after extensive optimisation, editing outcomes are often inefficient and heterogeneous(2–7). Consequently, variant-associated phenotypes are obscured unless correctly edited cells can be genotyped and enriched. Conventional workflows typically rely on single-cell isolation, clonal expansion and subsequent genotyping(8), but these approaches are time-consuming, inherently low-throughput, are limited to cells with sufficient potential for expansion and are prone to variability and bias due to single cell bottlenecking and expansion.

Single-cell RNA sequencing (scRNA-seq) coupled with genetic perturbation enables high-throughput interrogation of the effects of genetic variation on the cellular transcriptome(9, 10), but most methods use the guide RNA as a proxy of genotype, limiting its use to highly efficient editing methodologies such as CRISPR knockout, activation or interference (CRISPRa/i). Methods that couple whole genome sequencing with transcriptomic profiling within single cells have been developed(11–17). Although this approach provides genome-wide coverage, whole genome sequencing costs are very high, limiting scale, and these approaches additionally suffer from high dropout rates, limiting their utility in interrogating the function of specific variants.

Another class of methods detect variants directly within cDNA using short or long read sequencing(18–21), but these are limited to highly transcribed regions of genes, and are blind to variants that would result in loss of transcript abundance such as frameshift indels that result in nonsense mediated decay of the transcript or changes in splicing that splice around the variant.

In contrast, Target-seq(22) and CRAFT-seq(23) incorporate targeted amplification of genomic DNA alongside whole-transcriptome amplification, enabling accurate genotyping of selected loci. However, these approaches do not enable enrichment of edited cells prior to sequencing. Droplet microfluidics-based methods, such as scSNV-seq(6), combine targeted single-cell genotyping with parallel scRNA-seq through cell barcodes integrated in the genome. This offers increased scalability but requires upfront barcoding and specialized instrumentation.

All of these approaches lack the ability to prospectively identify and enrich SNP-defined cells before sequencing. As a result, when editing efficiency is low or heterogeneous, sequencing data is dominated by unedited or incorrectly edited cells, limiting scale and increasing experimental cost.

An effective approach would enable direct identification of SNP-defined single cells prior to sequencing while preserving full-length transcriptome information within a streamlined workflow. SNP-specific PCR strategies, including the use of phosphorothioate linkages, can achieve highly selective amplification of SNP alleles(24, 25). However, these approaches are typically limited to end-point genotyping and have not been integrated with whole-transcriptome sequencing workflows, which requires specific detection of the genotyping amplicon in the presence of other, full length cDNA amplicons.

Fluorescent detection strategies, such as TaqMan probes and molecular beacons, can detect specific amplicons but are relatively insensitive, and typically require locus-specific probe design and extensive optimization. In contrast, CRISPR-associated nucleases such as Cas12a provide programmable DNA recognition through simple redesign of guide RNAs (crRNAs), enabling flexible and scalable target detection(26, 27) and additional signal amplification by exploiting their “collateral” cleavage activity on fluorescent single-stranded DNA reporters. Cas12a-based platforms such as DETECTR(27) can discriminate single-nucleotide variants through guide RNA design but still requires target-specific optimization and is constrained by protospacer-adjacent motif (PAM) availability, limiting its general applicability as a standalone SNP detection strategy.

Here, we introduce SELECT-seq, a one-pot method that integrates highly SNP-specific PCR amplification with Cas12a-mediated fluorescence detection during whole-transcriptome amplification. This combines the benefits of specificity in the PCR and flexibility in the fluorescent detection. Importantly, SELECT-seq enables pre-sequencing identification and enrichment of SNP-bearing single cells without clonal expansion or additional genotyping steps, facilitating rapid functional interrogation without specialized instrumentation.

## Results

### SELECT-seq Enables Pre-Sequencing Enrichment of SNP Edits During Whole-Transcriptome Amplification

To enable pre-sequencing identification of SNP-bearing single cells, SELECT-seq integrates allele-specific PCR amplification with Cas12a-mediated fluorescence detection whilst simultaneously performing whole-transcriptome amplification from single cells (Fig. 1A). The workflow incorporates 3′ phosphorothioate-modified allele-specific primers, enabling selective amplification of a target locus without disrupting full-length cDNA generation and Cas12a to specifically detect the target locus amplicon within the mixture, allowing rapid identification of SNP-positive wells.

**Figure 1.**
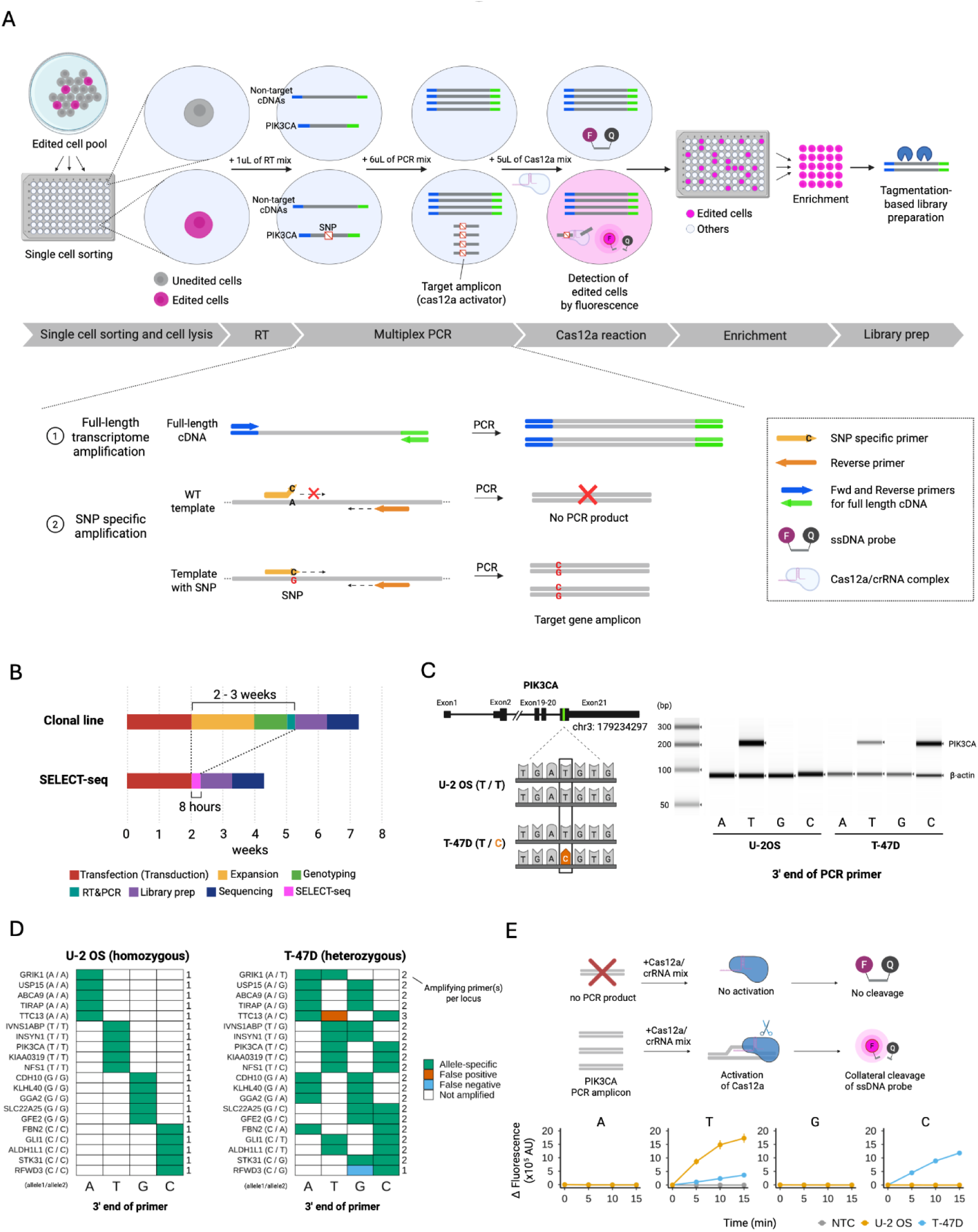
SELECT-seq enables pre-sequencing SNP detection during whole-transcriptome amplification. (A) Schematic of the SELECT-seq workflow. Single cell mRNA undergoes reverse transcription and whole-transcriptome amplification incorporating a 3′ phosphorothioate-modified SNP-specific primer. Cas12a-mediated fluorescence detection is performed directly on PCR products, enabling SNP identification prior to sequencing. (B) Timeline comparison of SELECT-seq and conventional clonal expansion workflows. (C) Validation of SNP-specific amplification at the *PIK3CA* locus using genomic DNA as input. Sequences are shown on the primer-binding (antisense) strand. The PIK3CA c.3463A>G variant is represented as the complementary T>C substitution. Four primers differing at the 3’ terminal nucleotide were tested in U-2 OS and T-47D. (D) SNP-specific amplification across 19 additional loci using U-2 OS (homozygous, left) and T-47D (heterozygous SNP, right) genomic DNA. (E) Schematic of Cas12a-mediated fluorescence detection following PCR amplification (top) and corresponding change in fluorescence (ΔF) across four *PIK3CA* PCR conditions (bottom).

Unlike previous strategies that physically separate transcriptomic amplification and genotyping, SELECT-seq operates as a true one-pot workflow. This minimizes sample loss, simplifies handling and automation, and is well suited for low-input and single-cell applications. Importantly, SELECT-seq allows SNP detection prior to sequencing, enabling targeted enrichment of specific genotypes. Whereas conventional workflows require single cell clonal expansion over two to three weeks before genotyping, SELECT-seq completes SNP detection and whole-transcriptome amplification within 8 hours (Fig. 1B).

We first validated SNP-specific amplification at the *PIK3CA* locus using genomic DNA as a template. Primers were designed to target a known PIK3CA mutation (NM_006218.4:c.3463A>G, chr3:179234297) present in the T-47D cell line (Fig. 1C, left panel) with the final base of each primer overlapping the SNP and phosphorthioate modifications on the final three positions. In U-2 OS cells, amplification occurred exclusively with the perfectly matched primer and not with the other three bases at this position, whereas in T-47D cells, amplification was observed with both T- and C-matched primers, consistent with the expected heterozygous SNP state (Fig. 1C, right panel). In some sequence contexts, even a single phosphorothioate bond was sufficient to inhibit amplification, although three consecutive bonds further reduced non-specific amplification (Fig. S1A). Also, even a mismatch at the second or third nucleotide from the 3′ end of the primer could effectively block amplification in some cases (Fig. S1B).

To assess generalizability, we used the COSMIC(28) database to identify 19 additional homozygous SNPs in U-2 OS cells that were heterozygous in the T-47D cell line. For each SNP we designed four primers that recognised all possible bases in that position. In U-2 OS, only the perfectly matched primer amplified each target, while in T-47D, two primers per gene typically amplified, reflecting the correct heterozygosity (Fig. 1D). The genotype was called correctly 98.7% of the time, with only one false positive and one false negative amplification across these 76 primer pairs across two cell lines. This confirms that phosphorthioate-modified primers can reliably discriminate alleles across diverse genomic contexts.

We next tested whether Cas12a-mediated fluorescence detection could be performed directly on PCR products. Cas12a activates robust collateral cleavage of fluorophore- and quencher-labelled single-stranded DNA (ssDNA) probes only in the presence of the target amplicon(27), producing a rapid and quantifiable fluorescence signal (Figure 1E, top panel). To enable direct detection after PCR amplification, we optimised reaction buffer conditions (Fig. S2A-E) and evaluated four crRNAs designed using DeepCpf1(29) (Fig. S2F, G). While all crRNAs were functional, crRNA4 produced the strongest fluorescence signal, consistent with its higher predicted activity score(29). When Cas12a reaction mix was added to PCR products generated from U-2 OS and T-47D inputs, robust fluorescence was observed only in the presence of correctly amplified *PIK3CA* amplicons (Fig. 1E, bottom panel), consistent with the gel-based validation.

Together, these results demonstrate the specificity of phosphorthioate-modified SNP-specific primers as a generalisable method for detecting SNPs and rapid fluorescent detection of the resulting PCR products using Cas12a.

### SNP-Based Distinction of U-2 OS and T-47D Cell Lines While Preserving Transcriptome Integrity

As proof of principle, we used SELECT-seq to profile the whole-transcriptome of U-2 OS (wild-type T/T) and T-47D (*PIK3CA* heterozygous T/C mutant) cells at single-cell resolution. Individual cells from each cell line were sorted into 96-well plates and subjected to reverse transcription, followed by PCR using an allele-specific primer targeting the mutation only present in T-47D. Cas12a reaction mix was then directly added without purification, and the fluorescence was monitored for 30 mins (Fig. 2A).

**Figure 2.**
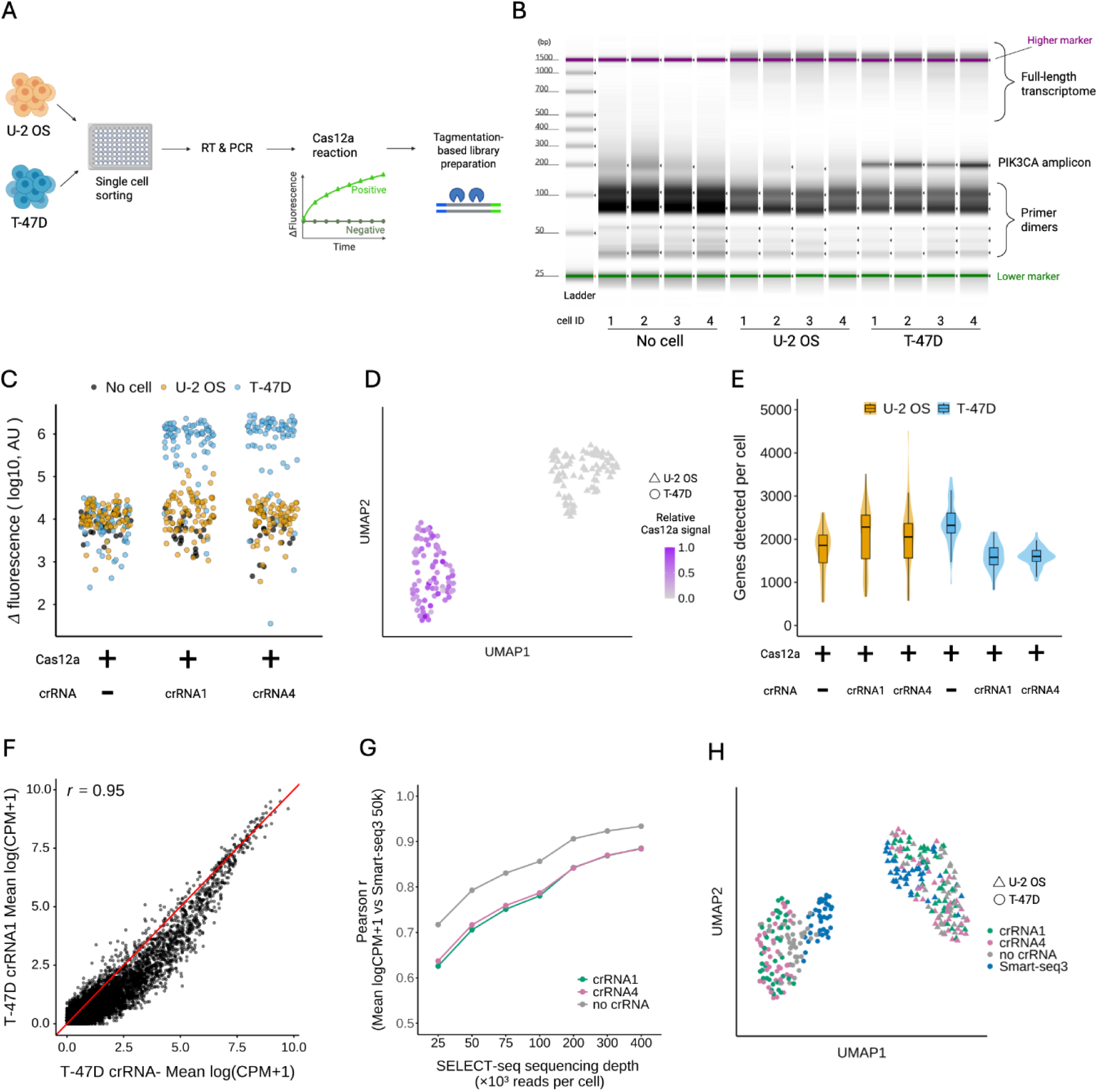
SELECT-seq distinguishes SNP-bearing single cells while preserving transcriptome integrity. (A) Schematic of the SELECT-seq workflow using U-2 OS (wild-type) and T-47D (*PIK3CA* mutant) cells. Cells were sorted individually into defined wells. (B) Gel electrophoresis analysis showing full-length cDNA and *PIK3CA* locus amplification from individual wells, with each lane representing a single cell or no-cell control. (C)(ΔF at the 10-minute time point following Cas12a-mediated fluorescence detection across individual wells. Values below zero are not displayed. (D) UMAP embedding showing correspondence between Cas12a signal and transcriptomic identity. (E) Violin plots of genes detected per cell, downsampled to 200, 000 reads per cell. (F) Pseudobulk expression correlation (mean logCPM+1) between crRNA− (n=40) and crRNA1 (n=42) conditions within T-47D cells. The red line indicates the identity line (y = x). (G) Pseudobulk expression correlation (mean logCPM+1) between Smart-seq3 (50, 000 reads per cell) and SELECT-seq across different sequencing depths. (H) UMAP embedding showing clear separation between U-2 OS and T-47D cells in Smart-seq3 (50, 000 reads per cell) and SELECT-seq (200, 000 reads per cell).

Gel electrophoresis confirmed co-amplification of the full-length cDNA (>500 bp) and the *PIK3CA* locus specifically in T-47D (Fig. 2B). As expected in U-2 OS, only whole-transcriptome amplification was observed with no SNP specific *PIK3CA* PCR product. No specific amplification products were observed in no-cell controls, although there was a weak product at a similar size. However, the Cas12a-dependent fluorescence readout clearly separated T-47D and U-2 OS cells, with negligible signal in no-cell wells (Fig. 2C). Cas12a signal also corresponded to transcriptomic identity, as shown by UMAP embedding (Fig. 2D), indicating accurate genotype discrimination at single-cell resolution.

We next assessed whether the Cas12a-mediated fluorescence detection step altered transcriptomic quality or complexity. In T-47D cells, where target locus amplification occurs, crRNA+ libraries exhibited a modest reduction in detected genes per cell relative to crRNA− libraries (Fig. 2E). In contrast, no reduction was observed in U-2 OS, suggesting this could be due to activity of the Cas12a enzyme. Despite this difference, pseudobulk expression profiles (mean logCPM+1) remained highly correlated between crRNA− and crRNA+ conditions within each cell line (Pearson r ≈ 0.95; Fig. 2F, Fig. S3A-E). This indicates minimal distortion of relative gene expression, most of which was observed at low expression levels. In addition, expression profiles between different crRNAs targeting the same locus were highly correlated (Pearson r = 0.97, Fig. S3B), indicating a lack of crRNA-specific transcriptomic perturbation.

To evaluate whether biological signals are preserved, we benchmarked SELECT-seq against a gold standard scRNA-seq method, Smart-seq3. At matched depth, SELECT-seq exhibited reduced sensitivity relative to Smart-seq3, as expected given the additional PCR cycles, co-amplification of a target-specific amplicon and Cas12a enzymatic reaction. However, correlation of pseudobulk expression profiles (mean logCPM+1) between U-2 OS and T-47D increased progressively with SELECT-seq sequencing depth, reaching Pearson r >0.85 (Fig. 2G). We did observe a modest reduction in correlation of expression levels in crRNA+ libraries compared to crRNA-(Fig 2G), but correlation of differential expression (U-2 OS vs T-47D log₂ fold change) was largely unaffected (Fig. S3F), indicating minimal impact of Cas12a-mediated fluorescence detection on biological contrast.

Consistent with this observation, marker gene overlap (top 100 genes) between SELECT-seq and Smart-seq3 increased with sequencing depth and stabilized beyond ∼200k reads per cell (Fig. S3G). The transcriptional distinction between U-2 OS and T-47D was clearly preserved in both crRNA- and crRNA+ conditions, with SELECT-seq cells closely aligning with the corresponding Smart-seq3 populations in the UMAP embedding (Fig. 2H). Together, these results demonstrate that increased sequencing depth compensates for reduced molecular efficiency while maintaining biologically meaningful differences between cell types.

### SELECT-seq enables direct enrichment of rare edited cells from pooled populations

To assess practical applicability, we next applied SELECT-seq to detect NRF2 T80K mutant cells from a pooled CRISPR prime-edited HAP1 population containing approximately 6.7% mutant cells (Fig. 3A, Fig S4A). The T80K substitution disrupts KEAP1 binding to NRF2, preventing ubiquitin-mediated degradation and resulting in activation of the NRF2 transcriptional programme(30). Using allele-specific and WT-specific primers, we separately identified and enriched T80K mutant and unedited cells based on Cas12a fluorescence.

**Figure 3.**
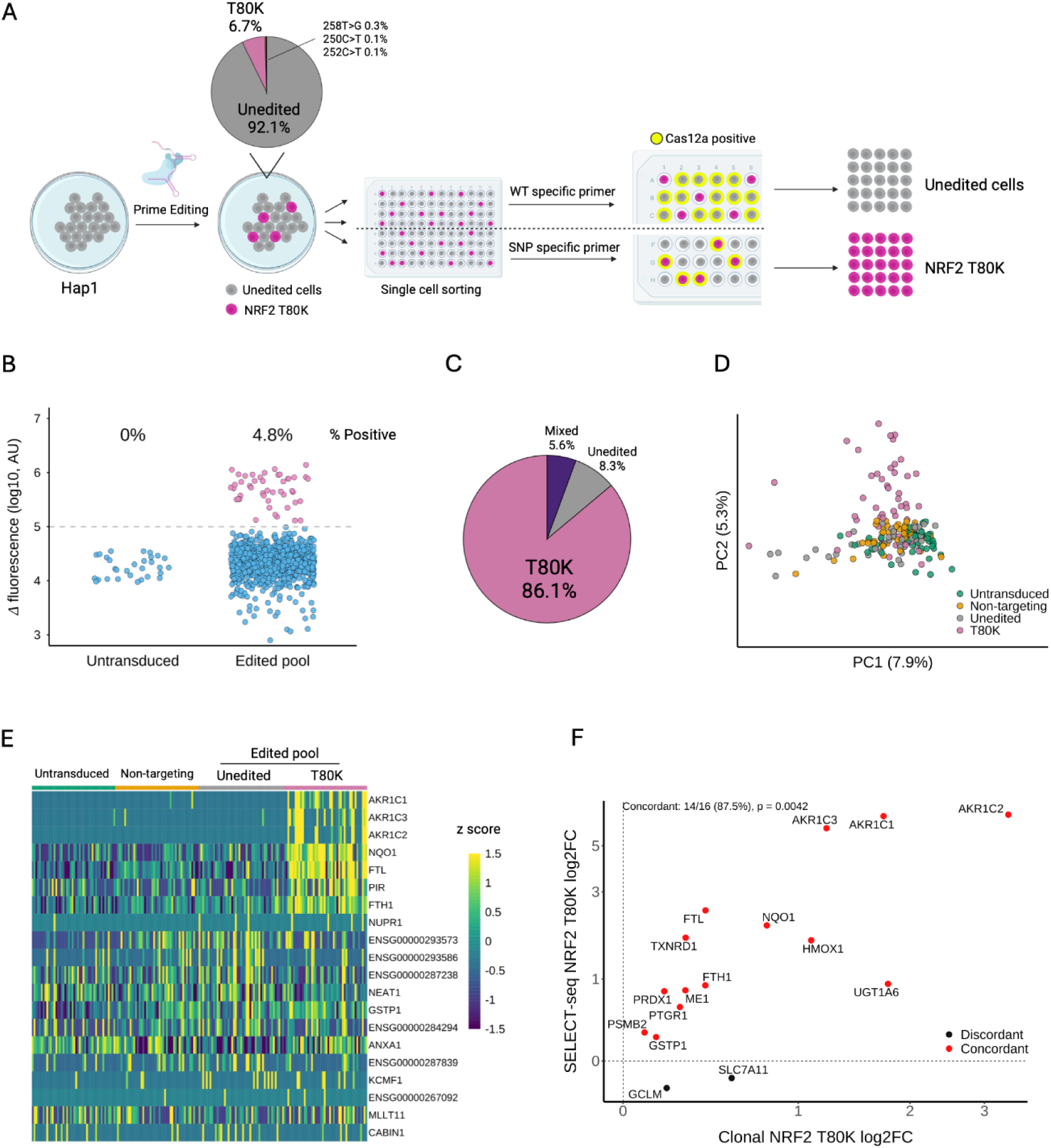
SELECT-seq enables enrichment and transcriptional profiling of rare edited cells from pooled populations. (A) Experimental design for detection of NRF2 T80K mutant cells from a pooled CRISPR-edited population. WT-specific or SNP-specific primers were used separately to detect unedited or NRF2 T80K cells, respectively. (B) ΔF at the 30-minute time point following Cas12a-mediated fluorescence detection across individual wells. A fluorescence threshold of 100, 000 was used to distinguish positive and negative populations, with 4.8% (46/960) of wells classified as T80K-positive. Values below zero are not displayed. (C) Genotype validation of fluorescence-gated cells demonstrating 86% (31/36) genotype accuracy. “Mixed” represents cells with ambiguous genotypes. (D) Principal component analysis of the top 100 highly variable genes across indicated conditions. Each point represents a single cell, plotted by PC1 and PC2. (E) Heatmap showing scaled gene expression of top 20 PC2-loading genes, indicating activation of NRF2-associated transcriptional programme. (F) Concordance of differential expression between Cas12a-enriched NRF2 T80K cells and clonal NRF2 T80K cells. Each point represents a high-confidence NRF2-responsive gene, defined as an NRF2 core gene with FDR < 1 × 10⁻⁶ in the clonal NRF2 T80K dataset. Genes are coloured according to whether the direction of log₂ fold change is concordant between datasets. The percentage of concordant genes and the corresponding two-sided binomial test P value are indicated.

Cas12a-mediated fluorescence detection identified a distinct T80K-positive population, comprising 4.8% of wells (Fig. 3B), consistent with the expected mutant frequency. Genotyping of isolated cells confirmed high classification accuracy, with 86% (31/36) of T80K-gated cells carrying the correct genotype (Fig. 3C). Genotyping was performed using primers flanking the SNP, with one primer positioned outside the target PCR amplicon to minimize potential bias associated with SNP-specific primer binding (Fig. S4B). Misclassified cells clustered near the fluorescence threshold, indicating that a higher threshold would improve classification accuracy (Fig. S4C). Similarly, detection of the WT allele was highly accurate, with 94% (32/34) of cells confirmed as the correct genotype (Fig. S4B, C, E). These results demonstrate that SELECT-seq enables reliable identification and enrichment of rare, edited cells directly from pooled populations.

Transcriptomic profiling of enriched cells revealed a clear shift in principal component space relative to unedited controls (PC2; Fig. 3D). Among the top 20 genes contributing to this component, NRF2-related genes were strongly upregulated in Cas12a-enriched T80K cells (Fig. 3E), consistent with activation of the expected transcriptional programme. In contrast, no clear differences were observed across other principal components (PC3–5; Fig. S5A).

Differential expression analysis between Cas12a-enriched T80K and unedited cells showed strong agreement with an independently generated clonal T80K reference dataset. Among high-confidence NRF2-responsive genes, defined as NRF2 core genes with FDR < 1 × 10⁻⁶ in the clonal NRF2 T80K reference dataset, 87.5% (14/16) exhibited concordant directional changes between datasets (binomial test, *P* = 0.004; Fig. 3F) demonstrating the effectiveness of SELECT-seq in recapitulating the effects of this variant. Comparison of gene expression patterns with both the clonal NRF2 T80K and clonal KEAP1 knockout reference datasets showed that, although the overall transcriptional response was highly concordant, a subset of genes displayed discordant expression changes (Fig. S5B). These differences may reflect clonal adaptation, technical variability, or biological effects that differ between rapid enrichment of edited cells and long-term clonal expansion. Some of these discordant genes showed expression changes that were consistent with the clonal KEAP1 knockout reference dataset but not the clonal NRF2 T80K dataset, suggesting that at least some differences may represent genuine NRF2-dependent transcriptional responses that were not detected in one of the two T80K datasets. Notably, several established NRF2 target genes associated with pathogenic NRF2 activation, including AKR1C1/2/3, NQO1, and FTL, were consistently upregulated across all three datasets, supporting preservation of the canonical NRF2 transcriptional programme. Together, our results demonstrate that SELECT-seq captures biologically meaningful genotype–phenotype relationships without requiring clonal expansion.

## Discussion

Precise linkage of genetic variation to its effects on cellular transcriptome and function remains a central challenge in functional genomics. Here, we introduce SELECT-seq, which enables pre-sequencing identification and enrichment of SNP-edited single cells alongside whole-transcriptome amplification. It operates as a true one-pot workflow without requiring additional or parallel genotyping steps and enables profiling of low-frequency edited cells directly from pooled populations. These features make SELECT-seq particularly useful for efficient functional interrogation of specific variants without specialised instrumentation, including in primary or *ex vivo* cell types (e.g. T cells and haematopoietic stem cells) where clonal expansion is not feasible.

Despite these advantages, several limitations should be considered. First, the Cas12a fluorescence readout is binary and does not distinguish heterozygous from homozygous edits. Whilst this still allows enrichment of cells with correct edits, it means that the genotype of the selected cells needs to be sequenced in a subsequent step (Fig. 3C). Second, the requirement for increased sequencing depth to achieve comparable data quality to the gold standard RNA-seq (e.g. Smart-seq3(31)), although with recent decreases in the cost of high throughput sequencing, this is becoming less of a problem. It would also be possible to improve transcriptomic quality by increasing the number of cells assayed by limited clonal growth after single cell isolation. Finally, plate-based implementation currently limits throughput relative to droplet-based methods. However, recent methods using programmable microfluidics (e.g. https://acxel.com/ (32, 33)) or semi-permeable hydrogel compartments for cell encapsidation (e.g. https://www.atrandi.com (34), https://www.cellanome.com (35)) could be used to overcome this and enable scaling to hundreds of thousands of cells.

In summary, SELECT-seq provides a rapid and accessible approach for linking genetic variation to transcriptional outcomes at single-cell resolution. By enabling pre-sequencing enrichment of genotype-defined cells, it expands the toolkit for functional genomics and the systematic characterisation of disease-associated variants. Our methodology broadens the range of cell types amenable to functional variant analysis, including primary cells that cannot readily undergo clonal expansion. Beyond variant validation, the approach may facilitate studies of rare somatic mutations in primary tissues by enabling enrichment of genotype-defined cells before transcriptomic profiling.

## Methods

### Cell Culture

U-2 OS (https://cellmodelpassports.sanger.ac.uk/passports/SIDM01191), T-47D (https://cellmodelpassports.sanger.ac.uk/passports/SIDM00097) and HAP1 (https://horizondiscovery.com/en/engineered-cell-lines/products/human-hap1-knockout-cell-li nes) cells were cultured in IMDM (Iscove’s Modified Dulbecco’s Medium) (12440053, Gibco), supplemented with 10% fetal bovine serum (FBS). All cells were maintained at 37 °C with 5% CO₂. Cells were stained with the Fixable Viability Dye eFluor 780 (65-0865-14, Invitrogen) to exclude dead cells before sorting into 96-well plates containing lysis buffer. HAP1 cells were co-stained with Hoechst 33342 (H3570, Invitrogen) to remove diploid cells.

### CRISPR Prime Editing

The HAP1 MLH1Δ cell line was obtained from the Leo Parts group (Wellcome Sanger Institute) and was generated as described(36). Briefly, HAP1 MLH1Δ cells (purchased from Horizon Discovery) were co-transfected with one plasmid expressing a PiggyBac transposase (pCMV-hyPBase55(36, 37)) and another plasmid containing a tetracycline-inducible PE2-P2A-GFP construct flanked by PB repeats (pPB-TREG3G-PE2-rtTA3G-P2A-eGFP(36)Addgene #196971). Cells were sorted based on GFP fluorescence and two clones were characterised for PE2 expression.

The pegRNA for generating the NRF2 T80K point mutation was designed using the Pridict online prime editing prediction tool (https://www.pridict.it/ (38)). The pegRNA sequences are listed in Supplementary Table 1. Sequences required for cloning the pegRNA into the chosen lentiviral vector (pKLV2-U6gRNA5(BbsI)-PGKpuro2ABFP-W(39) Addgene #67974) were obtained as gBlocks from IDT. Briefly, the vector backbone was linearised with BbsI and BamHI followed by gel extraction and subsequent PCR purification with Qiagen Qiaquick columns. The pegRNA insert was cloned into the prepared vector backbone by Gibson assembly and the resulting product was grown in overnight culture and plasmid purified. The completed vector was used to generate a stock of lentiviral particles by standard methods. Viral titre was determined by BFP fluorescence with the aim of identifying a low MOI to ensure that each HAP1 cell is infected by only a single viral particle.

For prime editing experiments HAP1 PE2 cells were seeded at 800, 000 cells per well of a 6-well plate the day prior to viral transduction (Day 0). Based on previous experiments determining titre, 20ul of viral supernatant equating to an MOI of between 0.2 and 0.3 was added to the culture medium on Day 1. In addition, polybrene was added at a concentration of 8µg/ml to aid transduction. Growth medium was changed on Day 2 to remove the virus and supplemented with 1µg/ml puromycin on Day 3 to select for successfully transduced cells. On Day 6 the medium was changed, removing puromycin and adding 2ug/ml Doxycyclin to induce expression of the prime editor. Cells were maintained in doxycycline until the end of the experiment at Day 17 at which point cells were harvested for genomic DNA extraction.

### Clonal line generation and characterization

NRF2-T80K HAP1 cells were created as described in Strauss et al. 2026 (40). Briefly, 1 µg plasmid expressing sgRNA (GTTACAACTAGATGAAGAGAC, in pMin-U6-ccdb-hPGK-puro(41)) and 2 µg homology directed repair donor templates (in the pMin backbone(41)) were transfected with 1.8 µl Xfect into HAP1-A5 cells constitutively expressing Cas9. On days 1 and 2 post-transfection, selection was performed with blasticidin (10 µg ml⁻¹) and puromycin (3 µg ml⁻¹; InvivoGen) to maintain Cas9 expression and select for transfected sgRNA plasmids. Cells were expanded over a further 10 days, stained with Hoescht (to select for haploid cells) and propidium iodide (to remove dead cells) and single, live, haploid cells sorted into 96 well plates. Clones were expanded and their genotype validated by PCR and Sanger sequencing.

### SNP-specific primer design

SNP-specific primers were designed using NCBI Primer-BLAST and synthesised by Integrated DNA Technologies (IDT) with standard desalting purification. Either the forward or reverse primer was constrained such that its 3′ terminal nucleotide matched the SNP of interest. Primer pairs with no or minimal predicted non-specific amplicons were selected for subsequent experiments. To enhance specificity, primers were modified with three consecutive phosphorothioate (PS) bonds at the 3′ end.

### Validation of SNP-specific primers

PCR was performed using KAPA HiFi HotStart Ready Mix (KK2601, Roche), with final concentrations of 0.2 μM each of forward and reverse primers targeting the gene of interest, 0.2 μM each of forward and reverse primers for ACTB (β-actin) as an internal control, and 0.2 ng μL⁻¹ genomic DNA (gDNA). Thermal cycling was performed as follows: 95 °C for 3 min; 30 cycles of 98 °C for 20 s, 65 °C for 15 s, and 72 °C for 20 s; followed by 72 °C for 1 min. PCR products were analysed on a TapeStation system (Agilent) by loading 1 μL of unpurified PCR product per sample. The concentration of target-gene amplicons was quantified when detected; undetected amplicons were assigned a value of 0. Values were first normalised to the ACTB amplicon and subsequently scaled to the maximum value within each gene and cell line. Values below 0.01 were considered not amplified. All primers used are listed in Supplementary Table 1.

### Cas12a reaction optimization

Reaction conditions were optimised by varying buffer composition, Cas12a concentration, crRNA concentration, and reporter probe design (Fig. S2). PCR conditions described in “Validation of SNP-specific primers” were used for optimisation, as the same polymerase and similar buffer compositions were used in SELECT-seq.

For optimization experiments, 10 μL of PCR product was directly mixed with 10 μL of Cas12a reaction mix, and fluorescence was measured at 37 °C for 30 min using a qPCR instrument. The change in fluorescence (ΔF) was calculated as the fluorescence at the indicated time point minus the fluorescence at 0 min.

### SELECT-seq workflow

#### Single-Cell Lysis and Reverse transcription

Single-cell lysis and reverse transcription were performed following the Smart-seq3 v3 protocol (https://dx.doi.org/10.17504/protocols.io.bcq4ivyw at protocols.io).

#### Multiplex PCR

Multiplex PCR was performed based on the Smart-seq3 v3 protocol, with minor modifications. The concentration of the forward and reverse primers for the whole-transcriptome amplification was reduced to 0.05 μM each (final). In addition, SNP-specific forward and reverse primers were included at a final concentration of 0.1 μM each. PCR was carried out using the same thermal cycling conditions as the Smart-seq3 v3 protocol, with the number of cycles increased to 35 to ensure sufficient amplification of SNP-targeted products.

#### Cas12a-mediated fluorescence detection

Following multiplex PCR amplification, 5 μL of Cas12a reaction mix was added directly to the PCR products without purification. The final reaction contained 50 nM Cas12a enzyme (M0653, NEB), 100 nM crRNA (IDT, desalted), 1 μM double-quenched FAM_ZEN_IBFQ ssDNA reporter (IDT, HPLC purified), and 1× NEBuffer r2.1. Fluorescence was monitored at 37 °C for 30 min using a qPCR instrument, followed by incubation at 65 °C for 10 min to inactivate Cas12a. The reaction was then stored at −20 °C until subsequent purification. All crRNAs and reporter probe sequences are listed in Supplementary Table 1.

ΔF was calculated as the fluorescence at the indicated time point minus the fluorescence at 0 min. A fluorescence threshold was defined based on the separation between negative and positive populations and used to classify wells as SNP-positive or negative.

#### Purification and library preparation

Following Cas12a-mediated fluorescence detection, 10 μL of reaction per well was mixed with 30 μL of nuclease-free water and 24 μL of AMPure XP beads (A63882, Beckman Coulter). Purification was performed according to the Smart-seq3 v3 protocol. DNA concentration was measured using the Qubit dsDNA HS Assay Kit (Q32851, Invitrogen) in 96-well plates. Samples were diluted to 100 pg μL⁻¹, after which tagmentation and index PCR were performed according to the Smart-seq3 v3 protocol with minor modifications. Tagmentation was terminated by addition of 0.1% SDS. Index PCR was performed for 14 cycles to ensure sufficient material for subsequent sequencing.

#### Read alignment and gene expression analysis

Raw sequencing data were processed and downsampled using zUMIs v2.9.7(35) to obtain UMI counts. Reads were aligned to the human reference genome (GRCh38 primary assembly) using STAR (v2.7.11b)(36). Gene-level counts were generated using GENCODE v46 annotation. Only exonic reads were retained for downstream analysis. For benchmarking analyses, cells with fewer than 3, 000 UMIs or more than 30% mitochondrial reads were excluded from both SELECT-seq and Smart-seq3 datasets. Gene-count thresholds were not applied because transcriptome complexity was evaluated as a benchmarking metric. Because the target gene is directly amplified during SELECT-seq, its expression level may be artificially inflated. Therefore, the targeted gene was excluded from downstream expression-based analyses. Specifically, *PIK3CA* was excluded from analyses of the U-2 OS/T-47D dataset, whereas *NFE2L2* was excluded from analyses of the NRF2 T80K dataset. The parameters used for data processing with zUMIs are available as described in the Code Availability section.

### Genotyping of Cas12a-positive single cells

Genotyping of Cas12a-positive single cells was performed by a three-step PCR strategy. In the first PCR, primers flanking the SNP were used, with the forward primer positioned outside the SNP-specific primer to minimise potential bias arising from SNP-specific primer binding. Purified SELECT-seq whole-transcriptome library (1 μL) was used as input for each 25 μL reaction. Each reaction contained 1× KAPA HiFi HotStart Ready Mix (KK2601, Roche) and 0.16 μM each of the forward primer (5′-cgacggaaagagtatgagctgg-3′) and reverse primer (5′-gcaacctgggagtagttggc-3′). Thermal cycling was performed as follows: 95 °C for 3 min; 35 cycles of 98 °C for 20 s, 65 °C for 15 s, and 72 °C for 20 s; followed by 72 °C for 1 min. Amplification was successful in ∼80% of cells. The concentration of the target amplicon was measured using a TapeStation system (Agilent).

A second PCR was performed to attach Nextera adapter sequences. First-round PCR products were diluted, and 5 ng of target amplicon was used as input for each 25 μL reaction. When the amplicon concentration was below 10 pg μL⁻¹, 0.5 μL of the first PCR product was used directly. Each reaction contained 1× KAPA HiFi HotStart Ready Mix and 0.16 μM each of the forward primer (5′-GTCTCGTGGGCTCGGAGATGTGTATAAGAGACAGcgacggaaagagtatgagctgg-3′) and reverse primer (5′-TCGTCGGCAGCGTCAGATGTGTATAAGAGACAGgcaacctgggagtagttggc-3′). Uppercase sequences correspond to Nextera adapter sequences and lowercase sequences to gene-specific regions. Thermal cycling was performed as follows: 95 °C for 3 min; 8 cycles of 98 °C for 20 s, 63 °C for 15 s, and 72 °C for 20 s; followed by 72 °C for 1 min. The resulting PCR product was then diluted fourfold for the subsequent indexing PCR.

For index PCR, 1 μL of the diluted second-round PCR product was used in each 25 μL reaction containing 1× KAPA HiFi HotStart Ready Mix and 0.16 μM each of Nextera i5 and i7 index primers. Thermal cycling conditions were: 95 °C for 3 min; 10 cycles of 98 °C for 20 s, 63 °C for 15 s, and 72 °C for 20 s; followed by 72 °C for 1 min.

Paired-end reads were merged using FLASH v1.2.11(42), and the region spanning the SNP ±20 bp was extracted. For each cell, a genotype was assigned as Unedited or NRF2 T80K when the most abundant allele accounted for at least 95% of reads. Cells not meeting this threshold were classified as Mixed.

### Genotyping of pooled edited cells

Genotyping of CRISPR prime-edited cell populations was performed to determine the relative abundance of each genotype. A first PCR was carried out using 160 ng of extracted genomic DNA in a 100 μL reaction. Each reaction contained 1× KAPA HiFi HotStart Ready Mix (KK2601, Roche) and 0.3 μM each of the forward primer (5′-TCGTCGGCAGCGTCAGATGTGTATAAGAGACAGcgacggaaagagtatgagctgg-3′) and reverse primer (5′-GTCTCGTGGGCTCGGAGATGTGTATAAGAGACAGactctgtacctgggagtagttgg-3′). Uppercase sequences correspond to Nextera adapter sequences, and lowercase sequences represent gene-specific regions. Thermal cycling conditions were as follows: 95 °C for 3 min; 25 cycles of 98 °C for 20 s, 65 °C for 15 s, and 72 °C for 20 s; followed by a final extension at 72 °C for 1 min. The concentration of the target amplicon was measured using a TapeStation system (Agilent).

For index PCR, 10 pg of the target amplicon from the first PCR was used in a 25 μL reaction containing 1× KAPA HiFi HotStart Ready Mix and 0.16 μM each of Nextera i5 and i7 index primers. Thermal cycling conditions were: 95 °C for 3 min; 10 cycles of 98 °C for 20 s, 63 °C for 15 s, and 72 °C for 20 s; followed by a final extension at 72 °C for 1 min.

Paired-end reads were merged using FLASH(42), and the region spanning the SNP ±20 bp was extracted. Genotype frequencies were calculated based on the relative abundance of each sequence.

### Smart-seq3 library preparation and sequencing

Smart-seq3(31) libraries were generated according to the Smart-seq3 v3 protocol(43) using 25 cycles of PCR amplification. Subsequent purification and library preparation steps were performed as described for SELECT-seq to enable a direct comparison between methods.

### Library purification and sequencing

Indexed libraries were purified using AMPure XP beads and sequenced on the Element Aviti platform using paired-end 150-cycle sequencing.

### Concordance analysis

For comparisons between crRNA− and crRNA+ libraries, SELECT-seq data were downsampled to 200, 000 reads per cell. For comparisons with Smart-seq3, data were downsampled to 50, 000 reads per cell. These depths were selected to maximize cell retention while maintaining sufficient transcriptome coverage for downstream analyses. For benchmarking analyses against Smart-seq3, SELECT-seq datasets were downsampled to the indicated sequencing depths, while Smart-seq3 was maintained at 50, 000 reads per cell.

To assess concordance of differential expression, pseudobulk expression profiles were generated by summing raw UMI counts across cells within each condition. Log₂ fold change values between U-2 OS and T-47D cells were calculated for each gene from pseudobulk counts using a pseudocount of 1. Pearson correlation coefficients were calculated between log₂ fold change values using genes shared between datasets.

For mean expression analysis, gene expression levels were normalised as counts per million (CPM) for each cell and log-transformed as log(CPM + 1). Mean expression values were then calculated across cells within each condition. Pearson correlation coefficients were calculated between mean expression profiles using genes shared between datasets.

For marker gene overlap analysis, marker genes were defined from pseudobulk log₂ fold change values as the top 100 genes with the highest positive log₂ fold change for U-2 OS or the lowest negative log₂ fold change for T-47D. Marker sets identified from SELECT-seq at each sequencing depth and crRNA condition were compared with those identified from Smart-seq3 data downsampled to 50, 000 reads per cell. Concordance was quantified using the Jaccard index, defined as the size of the intersection divided by the size of the union of the two marker sets.

### NRF2 gene set concordance analysis

To assess concordance of NRF2-associated transcriptional changes between clonal NRF2 T80K cells and prime-edited cells profiled using different platforms (10x Genomics and SELECT-seq), directional concordance was quantified. An NRF2 core gene set was defined based on Strauss et al. 2026(40). Differential expression data for the clonal NRF2 T80K cell line were generated by Strauss et al. 2026(40) and provided to the authors for downstream analyses. High-confidence NRF2-responsive genes were defined as NRF2 core genes with FDR < 1 × 10⁻⁶ in the clonal NRF2 T80K dataset. These genes were used for directional concordance analysis between the clonal NRF2 T80K and SELECT-seq datasets.

Differential expression analysis of SELECT-seq data was performed using the Wilcoxon rank-sum test implemented in Seurat (v5.1.0) (44). Only the log₂ fold change values from the SELECT-seq dataset were used for comparison with the clonal reference dataset. Genes with non-finite log₂ fold change values were excluded, and genes with zero log₂ fold change in both datasets were removed. Directional concordance was calculated as the proportion of genes with matching log₂ fold change directions between datasets. Statistical significance was evaluated using a two-sided binomial test against an expected concordance rate of 50%. For heatmap visualization, high-confidence NRF2-responsive genes were defined as NRF2 core genes with FDR < 1 × 10⁻⁶ in either the clonal NRF2 T80K or clonal KEAP1 knockout reference datasets.

### Dimensionality reduction

For visualization of transcriptional differences between U-2 OS and T-47D cells, SELECT-seq and Smart-seq3 datasets were downsampled to 200, 000 and 50, 000 reads per cell, respectively. These depths were selected to maximize cell retention while maintaining sufficient transcriptome coverage for downstream analyses. Highly variable genes were identified using Seurat with the variance-stabilizing transformation (VST) method, retaining the top 1, 000 or 2000 variable genes. Data were scaled, principal component analysis (PCA) was performed using the selected genes, and UMAP was generated using the first 10 principal components.

For analysis of NRF2 T80K cells, highly variable genes were identified using the VST method, retaining the top 100 variable genes. Scaled expression values were used for principal component analysis (PCA). For heatmap visualization, the top 20 genes with the largest absolute PC2 loadings were selected and displayed using scaled expression values.

## Author Contributions

S.I., Q.W, H.R., D.A., S.C. and A.B. conceived the work and designed experiments, T.B-S. made the clonal cell lines, S.I., D.G., H.R., T.B-S. and S.C. executed the experiments, S.I., A.W. and M.S. analysed data, S.I. and A.B. wrote the manuscript with input from all other authors.

## Declaration of Interests

A.B. and Q.W are founders of EnsoCell therapeutics for which A.B. is a consultant and Q.W. is now an employee. This work was part funded by Acxel.

## Data and code availability

Raw sequencing data have been deposited in the European Nucleotide Archive (ENA) under accession number PRJEB115314 and will be released upon publication. All scripts and parameters used for data processing are available on GitHub (https://github.com/ShoIwama/SELECT-seq).

## Funding

This work was funded by Acxel and Wellcome 220540/Z/20/A. For the purpose of Open Access, the author has applied a CC BY public copyright license to any Author Accepted Manuscript version arising from this submission.

## Acknowledgments

We thank the Cytometry Core Facility at Wellcome Sanger Institute (James Illing and Bee Ling Ng) for providing flow cytometry advice and cell sorting support. We acknowledge Scientific Operations at the Sanger Institute for support in next-generation sequencing. We thank members of the Bassett lab for helpful discussions and support. Schematics were created with https://BioRender.com. ChatGPT (OpenAI) was used to assist with language editing and code development. All results and conclusions were verified by the authors.

## Notes

https://github.com/ShoIwama/SELECT-seq

