## Supplementary Figures for "SELECT-seq allows Pre-Sequencing Enrichment of SNP Edits in One-Pot Single-Cell Whole-Transcriptome Sequencing"

### Figure S1

A

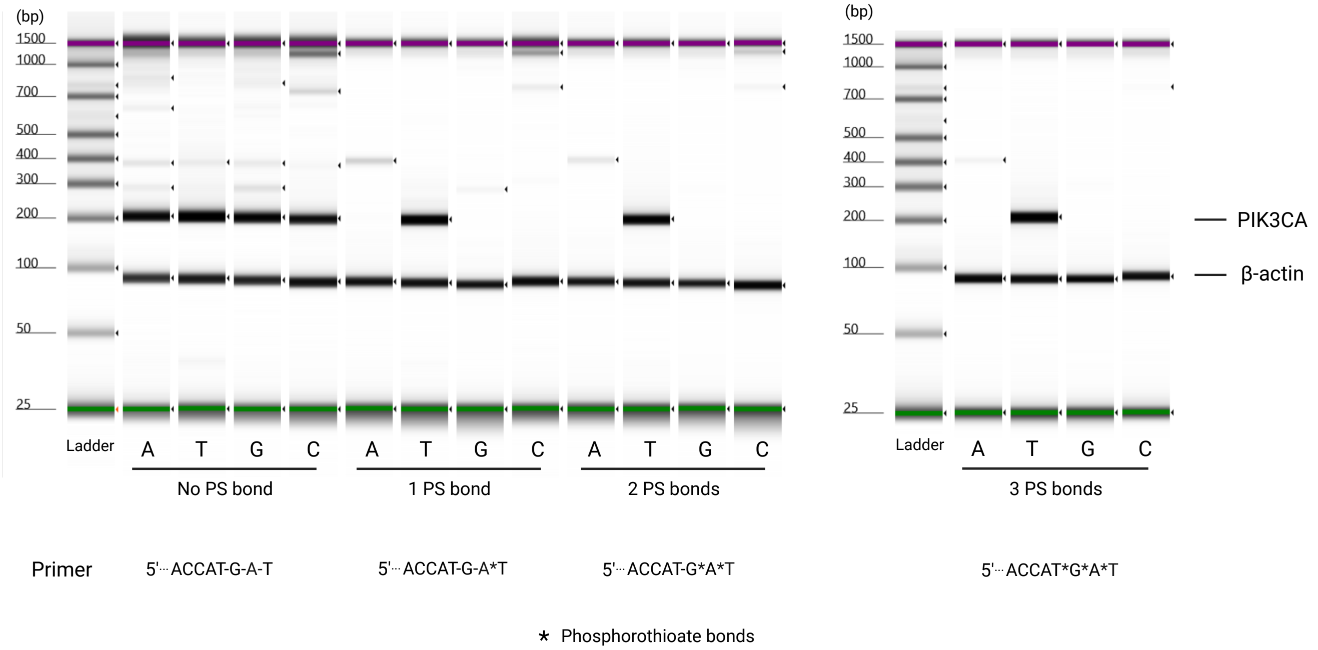


B

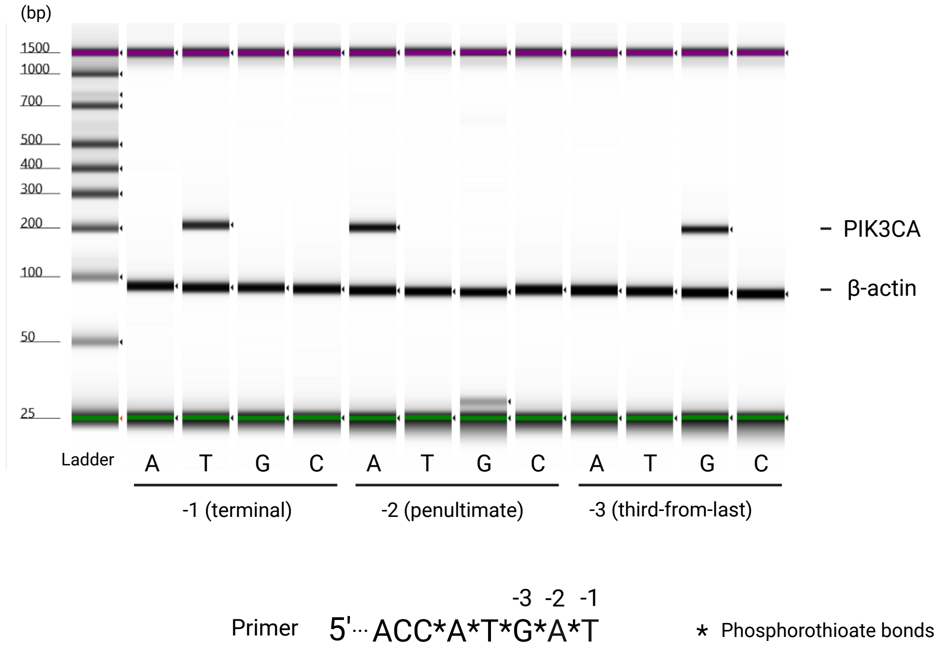


**Figure S1. Validation of phosphorothioate-modified primers and mismatch positioning for SNP-specific amplification.**
(A) Gel electrophoresis analysis of PIK3CA amplicons generated using primers containing zero, one, two, or three phosphorothioate (PS) bonds at the 3' end. Each lane corresponds to a primer differing at the terminal nucleotide. β-actin was amplified as a control.
(B) Gel electrophoresis analysis of PIK3CA amplicons generated using primers containing all possible nucleotides at the terminal (−1), penultimate (−2), or third-from-last (−3) position relative to the 3' end. Each base indicated corresponds to the nucleotide introduced at the specific position. β-actin was amplified as a control.

### Figure S2

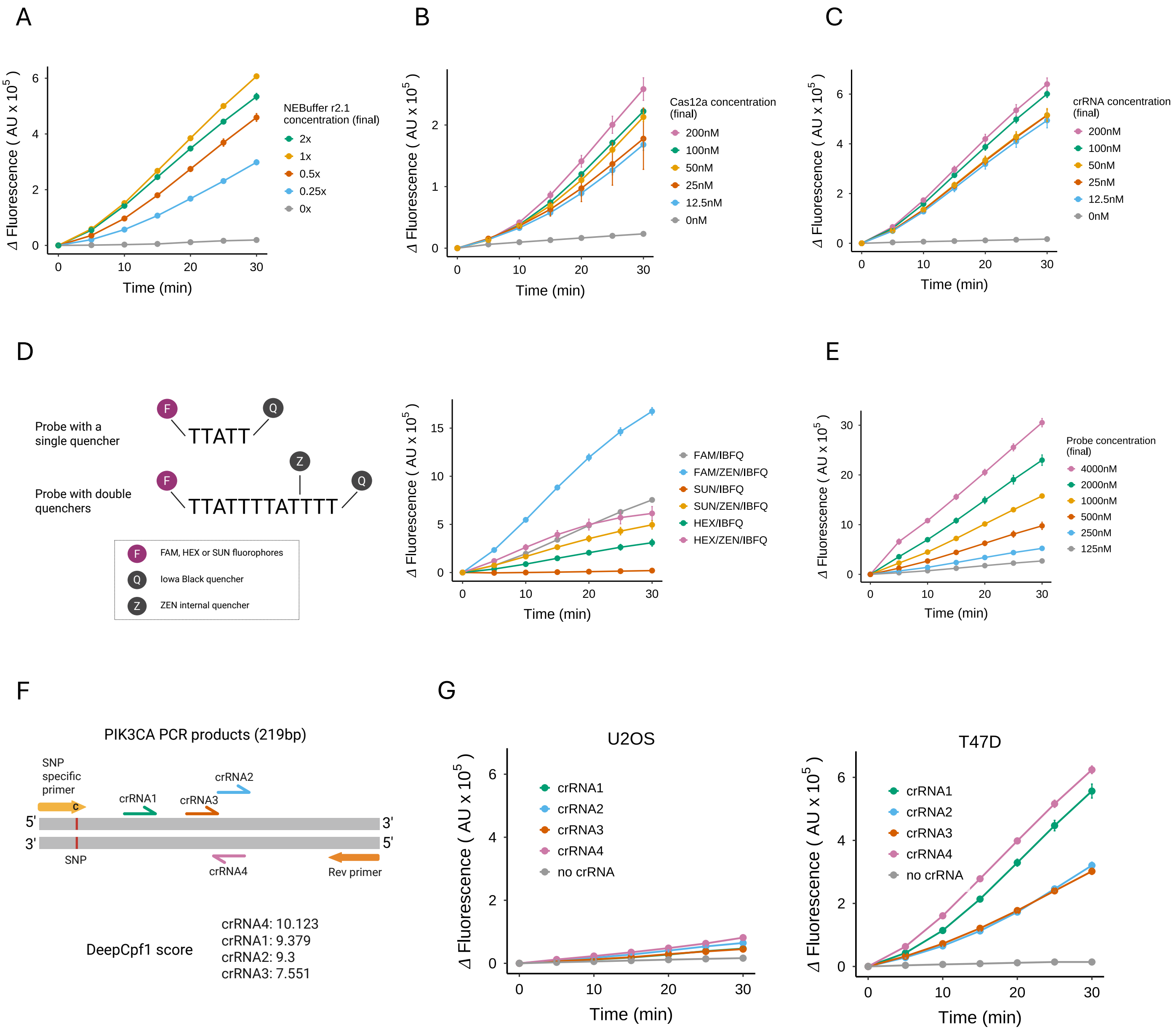


**Figure S2. Optimization of Cas12a reaction conditions and crRNA performance.**
(A-C) Δ fluorescence measured across varying final concentrations of NEBuffer r2.1 (A), Cas12a enzyme (B) and crRNA4 (C).
(D) Schematic illustrating the structures of reporter probes containing either a single quencher or double quenchers (left) and corresponding Δ fluorescence signals (right).
(E) Δ fluorescence across varying final concentrations of the FAM/ZEN/IBFQ reporter probe.
(F) Schematic showing the positions of the four crRNAs targeting the PIK3CA amplicon (top) and their predicted activity scores from DeepCpf1 (bottom).
(G) Δ fluorescence generated by crRNA1–4 or a no-crRNA control using PCR products from U-2 OS (left) and T-47D (right) templates.

### Figure S3

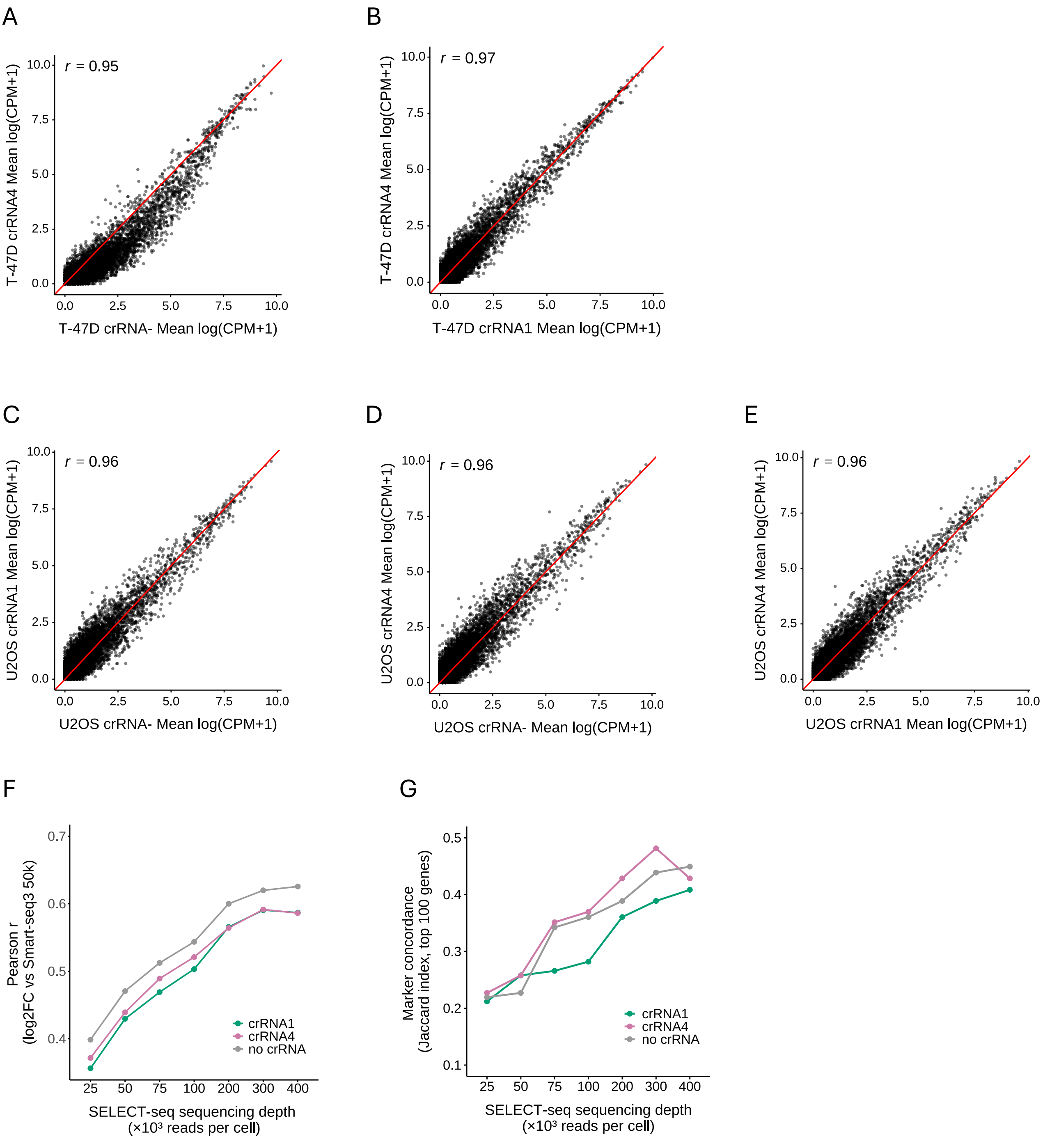


**Figure S3 Preservation of biological signals in SELECT-seq**

(A-B) Pseudobulk expression correlation (mean logCPM+1) between crRNA− and crRNA4 (A), and between crRNA1 and crRNA4 (B) in T-47D cells.

(C-E) Pseudobulk expression correlation between crRNA− and crRNA1 (C) , crRNA− and crRNA4 (D), and between crRNA1 and crRNA4 (E) in U-2 OS cells.

(F) Differential expression correlation (U-2 OS vs T-47D log2FC) between SELECT-seq and Smart-seq3 across sequencing depths.

(G) Marker concordance (Jaccard Index) based on the top 100 marker genes.

### Figure S4

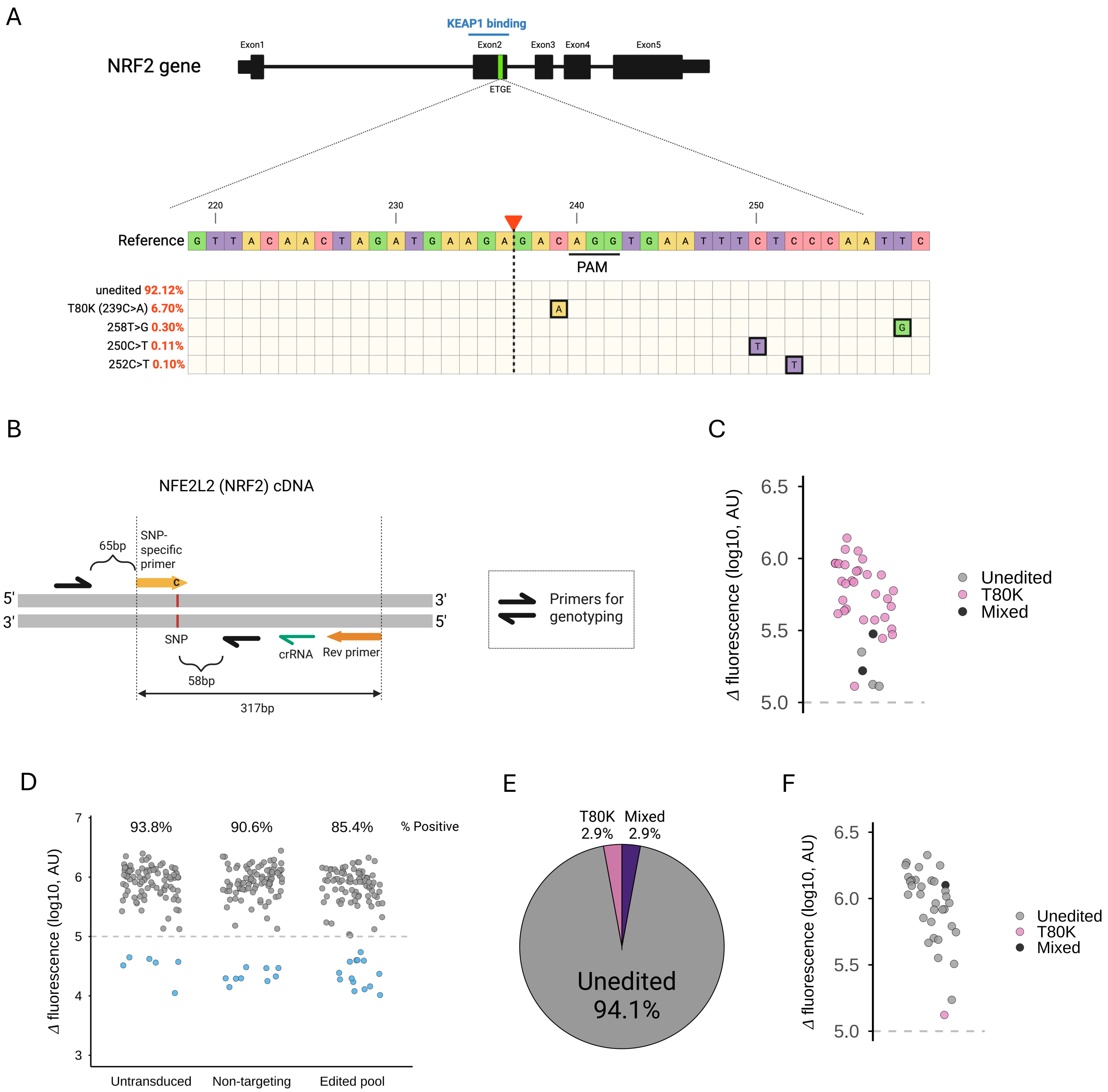


**Figure S4. Identification of NRF2 T80K and unedited cells from the pool of edited cells**

(A) Schematic showing the relative abundance of the top five mutations present in the CRISPR prime-edited cell pool.
(B) Schematic illustrating the positions of the primers used for genotype validation.
(C) Δ fluorescence distribution for successfully genotyped cells using the SNP-specific primer.
(D) Fluorescence-based identification of WT-positive wells.
(E) Genotype validation of fluorescence-gated unedited cells, demonstrating 94% positive predictive accuracy.
(F) Δ fluorescence distribution for the successfully genotyped cells using the WT-specific primer.

#### Figure S5

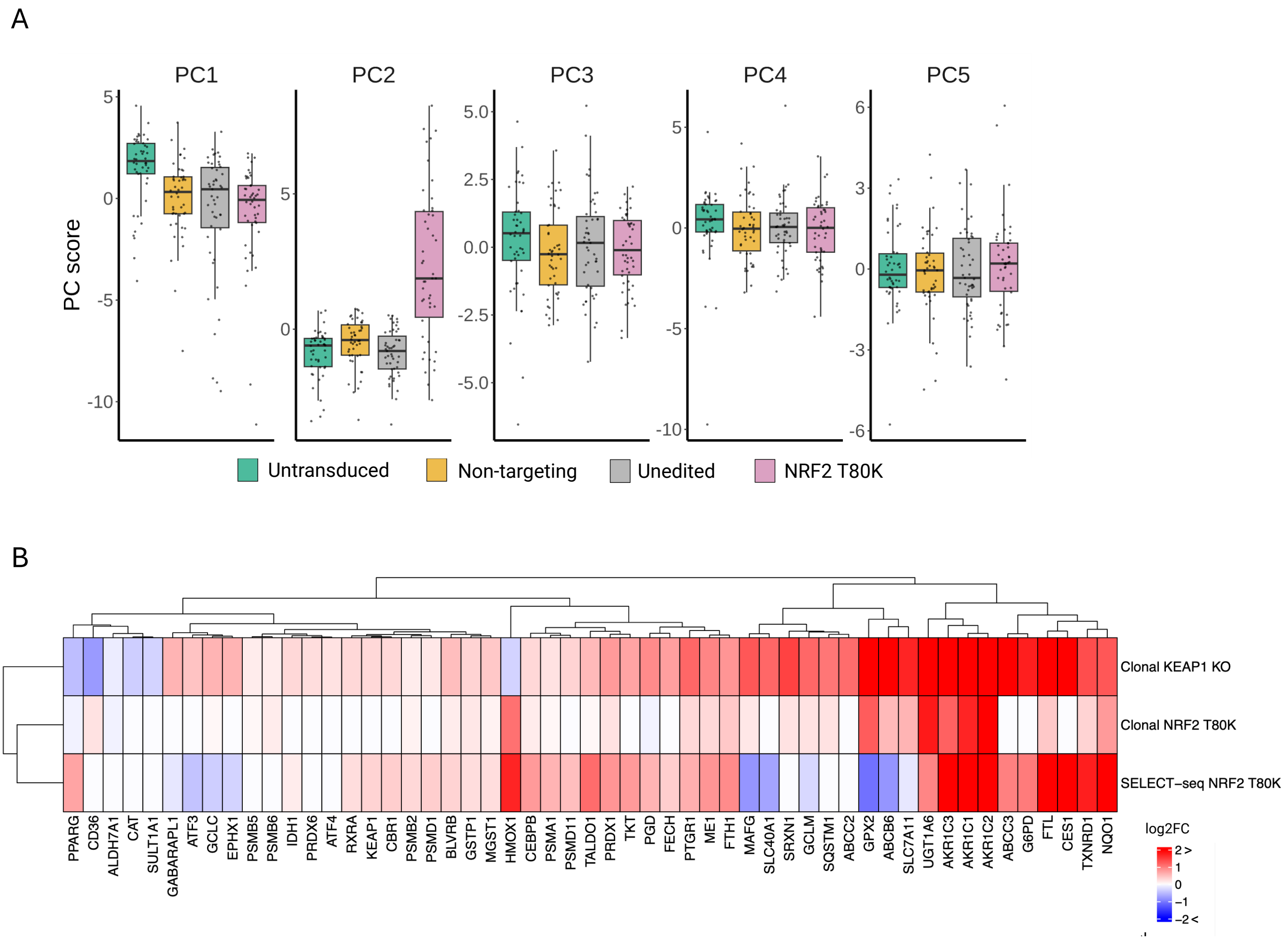


**Figure S5. Transcriptomic validation of Cas12a-enriched T80K cells.**
(A) Box plots showing the distribution of principal component scores (PC1–5) based on the top 100 highly variable genes across indicated conditions. (B) Heatmap of $\log_2$ fold changes for NRF2-associated genes, comparing Cas12a-enriched T80K cells with independently derived clonal T80K and KEAP1 KO lines. Only NRF2 core genes with FDR < $1 \times 10^{-6}$ in at least one of the clonal T80K or KEAP1 KO datasets were included. Values exceeding ±2 are capped at ±2.
